# Representing Sex in Cardiovascular Models: Calibrating Reference Parameters from Healthy Cohorts

**DOI:** 10.64898/2026.09.17.752488

**Authors:** Adhithi Lakshmikanthan, Felix Plappert, Weining Shen, Pim J.A. Oomen

## Abstract

Reduced-order models are increasingly used to study cardiac physiology and inform patient-specific therapies. However, a model’s prediction is only as reliable as its underlying parameters: representative model parameterization is essential to reflect the physiology of the populations these models are meant to represent, including biological sex. Most current models are parameterized from male or sex-agnostic data and/or focus on specific pathologies. Therefore, the goal of this work is to establish a formal parameter estimation pipeline for deriving reduced-order cardiovascular model parameter ranges that are physiologically representative of healthy women and men. We calibrated a closed-loop reduced order model of the heart and circulation separately for healthy female and male populations, using data pooled from eleven healthy cohorts. To account for parameter sensitivity and identifiability, we employed a three-stage parameter subset reduction pipeline: global sensitivity analysis (Sobol’s method), collinearity screening (Fisher information matrix), and profile-likelihood identifiability analysis. Sex-specific distributions of parameters that were deemed sensitive and identifiable for each sex, ten for women and 9 for men, were obtained by Hamiltonian Monte Carlo. All the calibrated parameters showed less than 80% overlap between sexes, with the smallest overlap observed in some of the most influential parameters, such as stressed blood volume. Comparing simulations of the calibrated models against allometrically size-matched simulations showed that body size explained some, but not all, of the sex differences. The resulting parameter distributions provide reference ranges usable in future mechanistic and patient-specific simulations to contribute to more inclusive cardiovascular modeling.

## 1 INTRODUCTION

The cardiovascular system is among the most studied physiological systems in the human body, yet its complexity continues to challenge our ability to fully characterize its behavior in health and disease. Computational models work synergistically with experimental and clinical studies by linking measurable data to the underlying mechanisms, enabling physiological function to be interpreted and predicted rather than only observed. Cardiovascular models are now used across scales and settings, from understanding physiological mechanisms to informing diagnosis and treatment planning in individual patients^1,2^. However, a model’s predictions are only as reliable as its underlying parameters: parameters that fail to represent the physiology of the individual or the population being studied could lead to inaccurate results. Representative model parameterization is therefore fundamental to cardiovascular modeling, so that they reflect the physiology of the people they are meant to represent, including biological sex.

Reduced-order models have been used extensively to study cardiovascular mechanisms and, increasingly, in precision-medicine approaches, because they retain essential physiological mechanisms governing cardiovascular function at low computational cost. Over the course of a century, these models have grown from Frank’s two-element Windkessel^3^ into full closed-loop representations of the heart and circulation. On the vascular side, the original arterial Windkessel was extended with characteristic impedance^4^ and inertance^5^ and embedded in closed-loop circuits partitioning the systemic and pulmonary beds into arterial and venous compartments^6,7^. On the cardiac side, prescribed pressure waveforms^6^ evolved into time-varying elastance^8^ and, ultimately, multi-segment formulations such as TriSeg that account for ventricular interaction through septal mechanics^9–11^. The result is a class of models rich enough to capture biventricular, closed-loop hemodynamics and now widely applied to both mechanistic and patient-specific questions^12–16^. However, the growing sophistication of reduced-order models of the cardiovascular system has led to a new challenge: each added component introduces new parameters that must be calibrated.

Recent work has addressed the model calibration challenge through formal parameter estimation. Sensitivity analysis and identifiability-guided subset reduction have been used to personalize reduced-order cardiovascular models to individual patients using deterministic approaches^17–20^, while Bayesian approaches now allow calibration against clinical data while quantifying parameter uncertainty^13,21–29^. These efforts, however, are almost exclusively disease-focused and/or patient-specific, and the reference values they build upon are predominantly derived from male subjects or non-sex-specific cohorts. This is a critical shortcoming, as sex differences are present throughout the cardiovascular system^30^, for example in cardiac chamber dimensions^31^ and vascular compliance^32,33^, with direct consequences for disease presentation and treatment outcomes. To date, the question of how reduced-order model parameters themselves differ between the sexes—and how much bias a sex-agnostic parameterization introduces—remains an open problem.

The goal of this work is to establish a formal parameter estimation pipeline for deriving reduced-order cardiovascular model parameter ranges that are physiologically representative of healthy women and men. We used our previously published model^13,34^ based on the TriSeg model to represent the heart and a closed-loop system consisting of five-element RCRCR compartments to represent the systemic and pulmonary arteries and veins. Following recent best-practice guidelines for mechanistic carpdiovascular model^21^, we employed a three-stage parameter subset reduction pipeline to identify sensitive and identifiable parameters: global sensitivity analysis (Sobol’s method), collinearity screening (Fisher information matrix), and profile-likelihood identifiability analysis. Parameter distributions representative of previously published healthy cohorts of women and men were obtained using Hamiltonian Monte Carlo with the No-U-Turn Sampler/ Differential Evolution Metropolis algorithm, propagating parameter uncertainty into all model outputs. The resulting parameter values accurately capture sex-specific trends in outputs held out from calibration, establishing healthy reference parameter ranges for women and men in reduced-order cardiovascular models that can inform future mechanistic and patient-specific studies.

## 2 METHODS

### 2.1 Target Data Collection for Sex-specific Calibration

Pressure and volume data for all four heart chambers were compiled from the literature to calibrate the biophysics model to a healthy female and a healthy male baseline state (Table 1). These values were obtained from 11 previous studies of healthy female and male subjects, with priority given to cohorts aged 50-65 years (Table 1). Reference values for right atrial mean pressure (RAMP), left atrial mean pressure (LAMP), mean pulmonary arterial pressure (MPAP) were only found for men, while right ventricular (RV) end-diastolic pressure (RVEDP) and left ventricular (LV) end-diastolic eccentricity index (LVEDEI) do not show sex-specific differences, and, LV differential pressure (LVdPSES) and RV differential pressure (RVdPSES) are assumed to be the same for both sexes. LVdPSES and RVdPSES represent the pressure increments corresponding to a 20% and 10% increase over the LV systolic pressure (LVSP) and RV systolic pressure (RVSP) respectively, modeling assumptions that are consistent with expected physiological pressure ranges.

**Table 1:** Sex-specific output targets for calibration. ^*′*^ marks outputs that show no sex-difference, while ^*⋆*^ marks outputs that are not sex-specific.

| Compartment | Output | Description | Age | n (Female) | Female | Male | Units | Ref |
| --- | --- | --- | --- | --- | --- | --- | --- | --- |
| Circulation | MAP | Mean Arterial Pressure | 60 ± 10 | 16 (50%) | 84.6 ± 6.67 | 88.6 ± 5.55 | mmHg | 35 |
|  | MPAP* | Mean Pulmonary Arterial Pressure | - | - | 15 ± 3.75 | 15 ± 3.75 | mmHg | 36 |
| Left ventricular (LV) | LVEDV | LV end-diastolic volume | 61 ± 9 | 852 (60%) | 109 ± 18 | 143 ± 24 | mm<sup>3</sup> | 37 |
|  | LVESV | LV end-systolic volume | 61 ± 9 | 852 (60%) | 35 ± 9 | 49 ± 12 | mm<sup>3</sup> | 37 |
|  | LVEDP | LV end-diastolic pressure | 49 ± 8 | 16 (50%) | 9 ± 2 | 13 ± 2 | mmHg | 38 |
|  | LVSP | LV systolic pressure | 49 ± 8 | 16 (50%) | 131 ± 5 | 133 ± 13 | mmHg | 38 |
|  | LVdPSES' | LV differential pressure | - | - | 15 ± 5 | 15 ± 5 | mmHg |  |
|  | LVEDEI' | LV end-diastolic eccentricity index | 36 ± 7 | 12 (-) | 1.01 ± 0.04 | 1.01 ± 0.04 | - | 39 |
| Right ventricular (RV) | RVEDV | RV end-diastolic volume | 51 – 60 | 51 (43%) | 127.8 ± 27.1 | 153.9 ± 19.2 | mm<sup>3</sup> | 40 |
|  | RVESV | RV end-systolic volume | 51 – 60 | 51 (43%) | 46.8 ± 13.1 | 64.6 ± 18.8 | mm<sup>3</sup> | 40 |
|  | RVEDP' | RV end-diastolic pressure | 18 – 88 | 18 (-) | 8 ± 2 | 8 ± 2 | mmHg | 41,42 |
|  | RVSP | RV systolic pressure | 18 – 90 | 507 (51%) | 24.7 ± 5.3 | 24.2 ± 5.6 | mmHg | 43 |
|  | RVdPSES' | RV differential pressure | - | - | 5 ± 2.5 | 5 ± 2.5 | mmHg |  |
| Left atrial (LA) | LAMaxV | LA max volume | 26 ± 4 | 434 (55%) | 89 ± 21 | 103 ± 30 | mm<sup>3</sup> | 44 |
|  | LAMinV | LA min volume | 26 ± 4 | 434 (55%) | 41 ± 11 | 46 ± 14 | mm<sup>3</sup> | 44 |
|  | LAMP* | LA mean pressure | - | - | 8 ± 2 | 8 ± 2 | mmHg | 36 |
| Right atrial (RA) | RAMaxV | RA max volume | 21 – 68 | 108 (42%) | 80 ± 17 | 108 ± 26 | mm<sup>3</sup> | 45 |
|  | RAMinV | RA min volume | 21 – 68 | 108 (42%) | 33 ± 11 | 50 ± 17 | mm<sup>3</sup> | 45 |
|  | RAMP* | RA mean pressure | 18 – 66 | 18 (-) | 4 ± 1 | 4 ± 1 | mmHg | 42,46 |

Note: Values are presented as mean ± SD. RVEDV and RVESV were converted from the reported mean and 5–95% confidence interval (CI) assuming normality. RVEDP, LAMP, RAMP and MPAP were reported without SD or CI; an SD of 25% of the reported mean was assumed.

### 2.2 Reduced-order model formulation

The reduced-order model consisted of two components: the heart and circulation. The heart was described using the TriSeg method^10^ (Fig. 1), with the ventricles modeled as three thick-walled spherical segments, the left free wall (LFW), right free wall (RFW), and septal wall (SW), interconnected at a circular wall junction. The left atrium (LA) and right atrium (RA) were each modeled as a single thick-walled sphere. No mechanical interaction between the atria and ventricles was assumed because of the dense connective tissue separating these chambers. The mechanics of each wall segment *W* ∈ { LFW, RFW, SW, LA, RA } were evaluated at its midwall surface, defined as the surface that divides the segment into two equal volumes. The segment volume was approximated by the product of its midwall reference area *A*_*m*,ref,*W*_ and its wall thickness *W*_th,*W*_ . Myofiber stretch was defined at the midwall with respect to the reference area, *λ*_*f,W*_ = *A*_*m,W*_ */A*_*m*,ref,*W*_ ^11^. Active myofiber stress *σ*_*f*_ was determined using a Hill-type model, scaled linearly by *S*_fact,v_ for the ventricles and *S*_fact,a_ for the atria^10^ whereas passive myofiber stress was determined using an exponential constitutive model, both as a function of *λ*_*f*_ ^34^. The three ventricular segments were mechanically coupled through equilibrium of their wall tensions, estimated from wall stress^11^, at the circular wall junction. At each time step, an iterative Newton scheme adjusted the segment curvatures until the normalized residual midwall tension at the junction dropped below 0.001.

**Figure 1:**
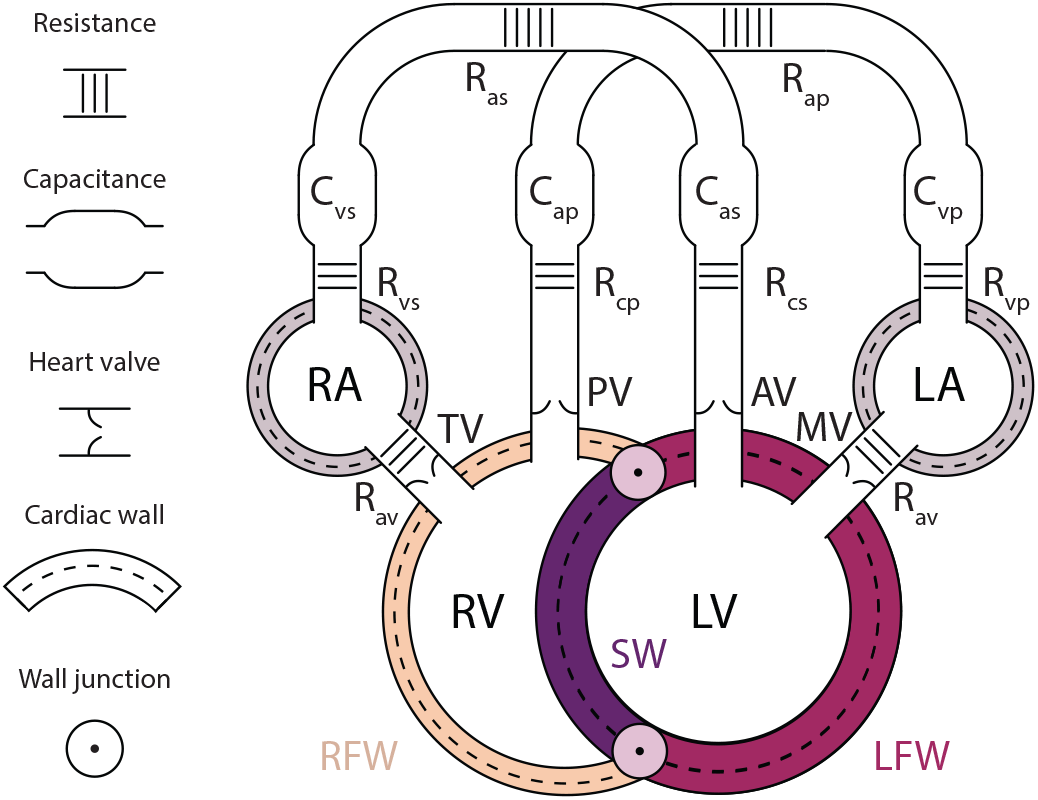
Schematic representation of the reduced order cardiac model. Cardiac mechanics is modeled using the Triseg formulation^10^, hemodynamics is modeled using the lumped-parameter model where systemic and pulmonary circulation is regulated using an electrical analog model of resistances, capacitances, and diodes. Figure adapted from^13^, licensed under CC BY 4.0.

The circulation was modeled using a lumped-parameter approach (Fig. 1). Each heart valve was modeled as pressure-sensitive diode, allowing blood to flow in one direction only, from the atria into the ventricles and from the ventricles into the arteries. The systemic and pulmonary circulations were each described by a five-element RCRCR Windkessel model. The total blood volume was functionally divided into a stressed blood volume (SBV) and an unstressed blood volume, the latter being the volume that the cardiovascular system can hold before its pressure exceeds 0 mmHg.

The ordinary differential equations of the heart and circulation were solved in a closed loop with a time step of 0.5 ms. At each step, the cardiac compartment pressures were computed from the current midwall tensions and curvatures. These pressures in turn set the midwall surface areas, from which the myofibre strain, stress and midwall tension were computed for the next iteration. The movement of blood between the heart chambers and the vasculature was computed using a forward Euler solver. Supplementary Section S1 describes in more detail how the TriSeg method solves the cardiac geometry and mechanics, and how the pressures and volumes are updated.

### 2.3 Parameter subset reduction

#### 2.3.1 Sensitivity analysis

A Sobol sensitivity analysis using the Saltelli sampling method^47^ and implemented in the SALib package^48^ was performed to identify the parameters that significantly contribute to the outputs’ variance. For a model output *Y* = *f* (***θ***), where *f* is the model (Section 2.2) and ***θ*** a vector containing all model parameters, the first-order Sobol index quantifies the fraction of output variance attributable to parameter *θ*_*j*_ alone,

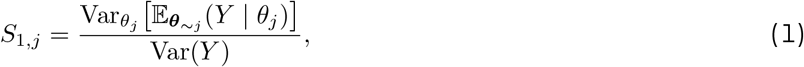

while the total-order index captures the contribution of *θ*_*j*_ including all interactions with other parameters,

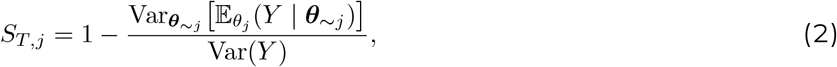

where ***θ***_*∼j*_ denotes all parameters except *θ*_*j*_.

Parameters were ranked by their total-order index *S*_*T,j*_ and the subset that cumulatively accounted for 90% of the total output variance was retained (Section 2.4). The remaining parameters were deemed non-influential and fixed at their nominal values or included in the latent space (Section 2.4).

#### 2.3.2 Identifiability screening

Parameters that passed the sensitivity screen were further assessed for identifiability due to collinearity using the Fisher Information Matrix (FIM)^49^. This step is complementary to the Sobol analysis: while *S*_*T*_ quantifies a parameter’s marginal contribution to output variance, it cannot detect collinearity between parameters whose effects on the output set are nearly parallel. The FIM is defined as

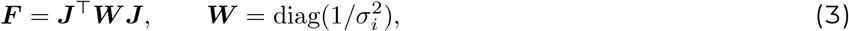

where ***J*** = *∂* ***ŷ*** */∂****θ*** is the Jacobian of the model outputs with respect to the parameters, evaluated at the midpoint of each parameter’s prior range, ***W*** weights each output by its inverse measurement variance 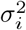, where *σ*_*i*_ is the population standard deviation of output *i* (Table 1). The Jacobian was estimated numerically using finite differences from the model evaluations already performed during the Sobol sampling, avoiding additional computational cost. Near-zero eigenvalues of ***F*** indicate unidentifiable directions in parameter space. Pairwise collinearity was assessed through the normalised column correlation |*C*_*ij*_ |of the weighted Jacobian; pairs exceeding |*C*_*ij*_ |≥0.8 were considered collinear, and one parameter from each pair was either fixed at its nominal value or moved to the latent space (Section 2.4). The parameter with the lower S1 value (Eq. 1) was fixed.

Profile likelihood (PL) analysis was performed for each remaining parameter to identify structurally and practically unidentifiable parameters^50^. This technique obtains a profile likelihood of each parameter *θ*_*j*_ by fixing it at a series of values and, at each fixed value, optimizing the remaining parameters to maximize the likelihood:

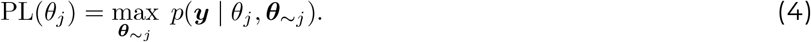

Each profile is constructed around the optimal estimate found by differential evolution^51^. Pointwise confidence intervals are defined by the range of *θ*_*j*_ values satisfying the threshold

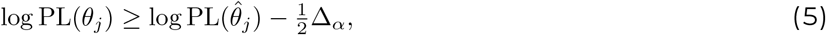

where 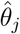 is the maximum likelihood estimate and 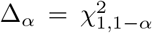 is the chi-squared quantile with one degree of freedom (e.g.,Δ_0.05_ = 3.84 for a 95% confidence interval). If its confidence interval has finite width within the prior range a parameter is considered practically identifiable. In contrast, if the profile is flat, the parameter is structurally non-identifiable; if the confidence interval extends to a prior bound it is practically non-identifiable–in either case they were then fixed at their nominal values or included in the latent space. We add end-diastolic stretch for the left free wall (*λ*_*ED, ℒ{⊒*_) and left atria (*λ*_*ED, ℒA*_) as additional targets during calibration from the PL step.

### 2.4 Model calibration

Bayesian inference is used to identify parameter distributions that are representative of the physiological variability of healthy male and female populations. The parameter vector ***θ*** = (***θ***^free^, ***θ***^lat^) comprises two groups. The free parameters ***θ***^free^, identified as influential and identifiable by the preceding sensitivity and identifiability analyses, were assigned independent uniform priors over physiologically plausible ranges (Table 2):

**Table 2:**
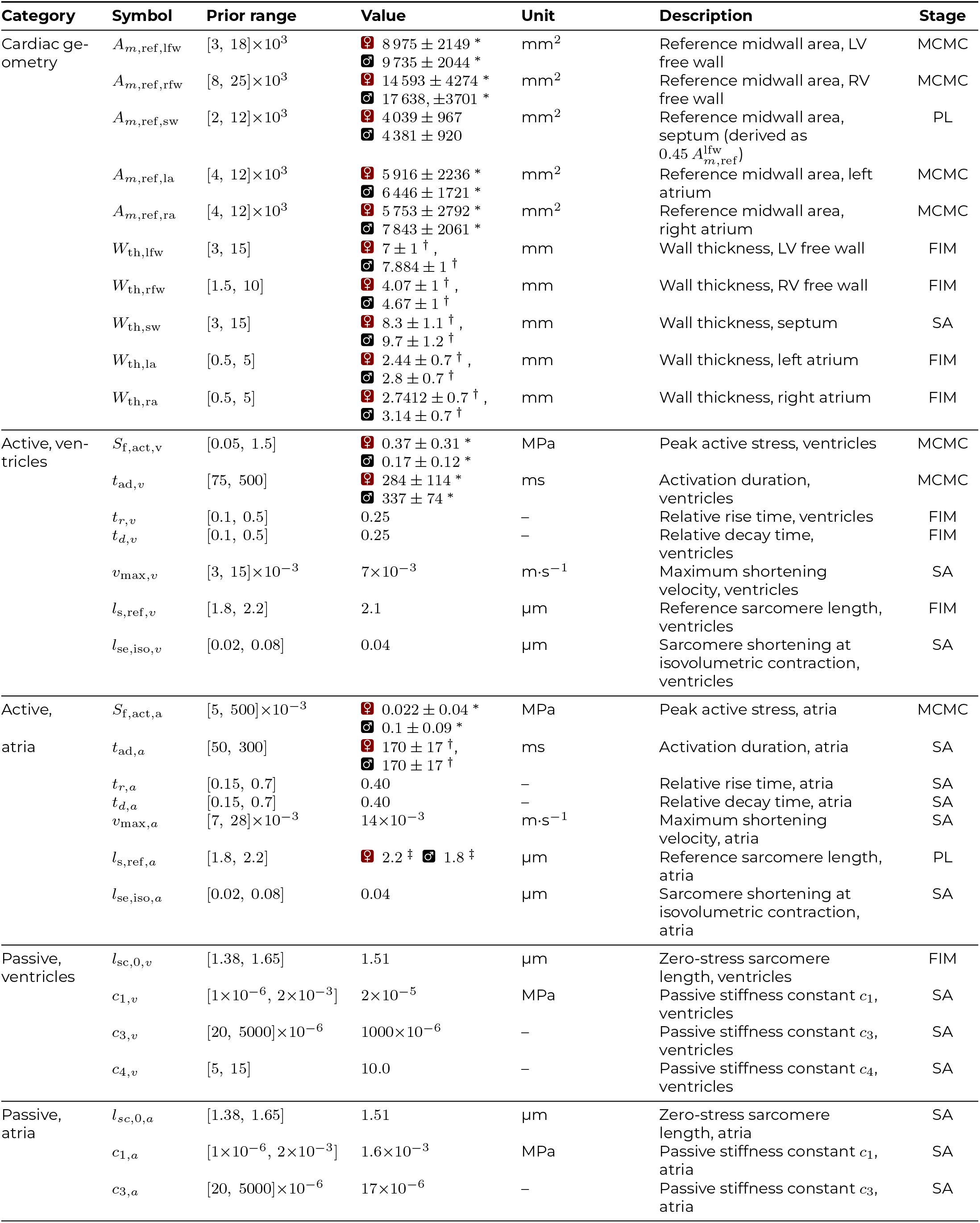
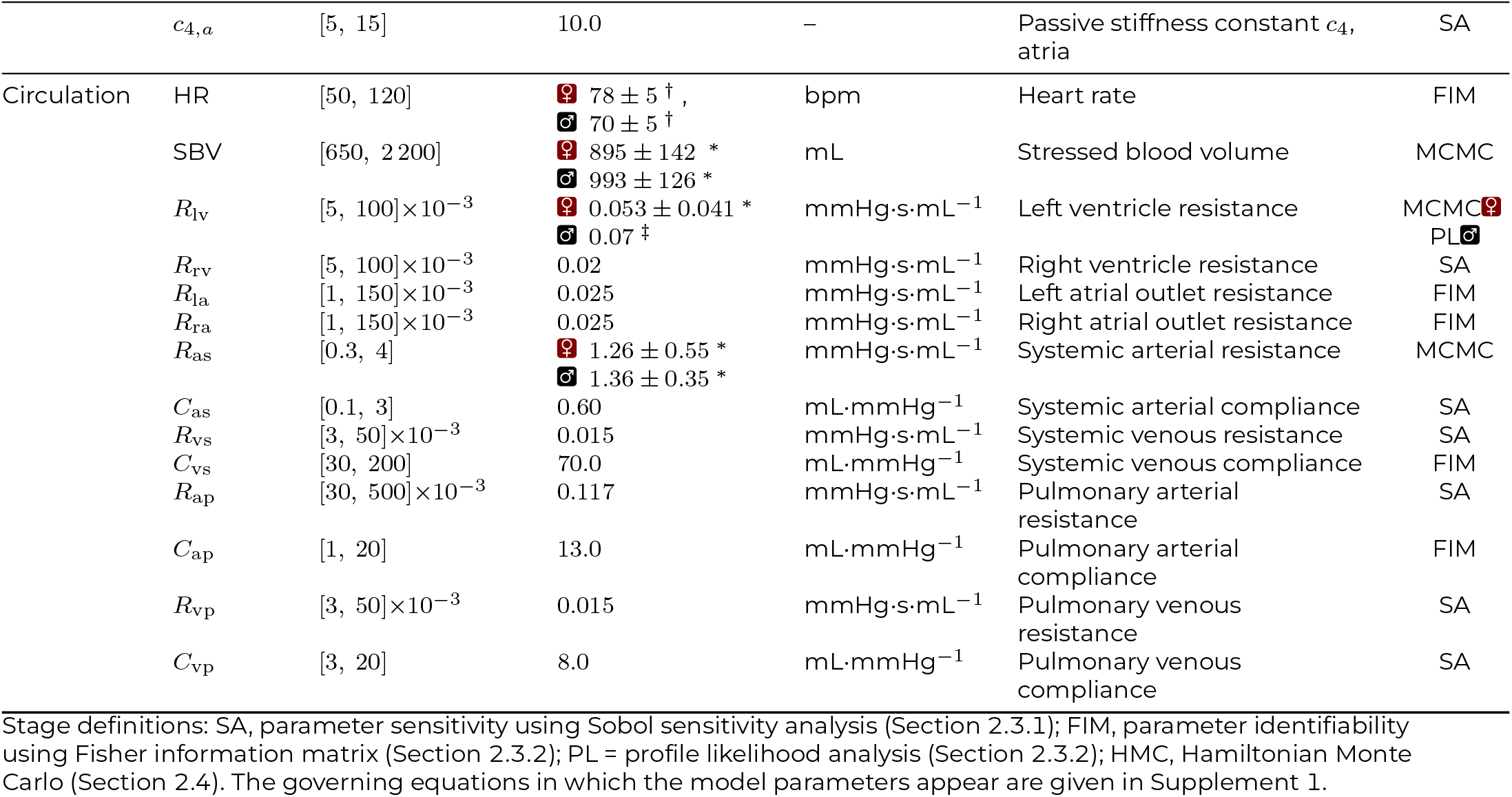
Model parameters with prior ranges and template starting values. Parameters marked ^*\**^ are inferred by Bayesian calibration using HMC with a uniform prior over the stated range; those marked ^*†*^ use an informed truncated-normal prior; those marked ^*‡*^ are set at PL optimized value; all others are fixed constants.

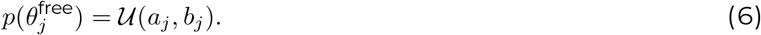

The latent parameters ***θ***^lat^ were obtainable from clinical measurements (e.g. end-diastolic wall thickness from echocardiography and heart rate from ECG). *W*_th,LFW_ and *W*_th,SW_ were obtained from Yeon et al.^37^ for both sexes. Where female data were unavailable (*W*_th,RFW_, *W*_th,LA_ and *W*_th,RA_), values were estimated as 12.7% lower than values reported in sex-unspecified or male-dominant studies^52,53^, consistent with the average sex difference observed in *W*_th,LFW_ and *W*_th,SW_. Rather than fixing these at nominal values, each was assigned a truncated normal 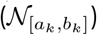 prior centered on its measured value, allowing their measurement uncertainty to propagate into the posterior:

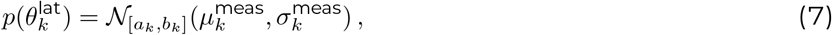

with bounds 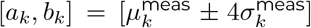 that serve as numerical guardrails against physiologically implausible proposals. To represent biological variability, each model output was assumed to follow an independent Gaussian likelihood centered around the sex-specific population mean:

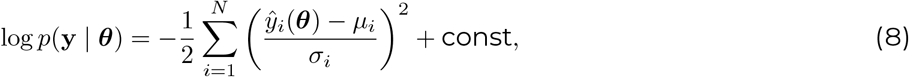

where *ŷ*_*i*_(***θ***) is the model prediction at parameter vector ***θ***, and *µ*_*i*_ and *σ*_*i*_ are the published population mean and standard deviation, resp., for output *i*. Note that assuming independent Gaussian distributions neglects correlations between outputs, however, because of the aggregated, population-level, data the output covariances required for a multivariate likelihood were not available.

The prior and posterior do not belong to the same distributions, and lack of conjugacy in the posterior necessitates the use of a Markov Chain Monte Carlo (MCMC) method as there is no closed-form solution. We adopt Hamiltonian Monte Carlo with the No-U-Turn Sampler (NUTS) as our primary sampler because it exploits analytical gradients available from the Gaussian process (GP) emulator to explore the smooth, correlated posterior far more efficiently than gradient-free alternatives such as random-walk Metropolis-Hastings^54^. Four independent chains were run, each drawing 1000 samples after 1000 warm-up steps during which the step size and mass matrix were adapted. Note that a substantial amount of additional samples are required to calculate the HMC gradients. Convergence was assessed by requiring the diagnostics 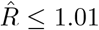 and bulk effective sample size (ESS_bulk_) *>* 400 for all parameters. Posterior predictive checks were performed by propagating the full set of posterior samples through the emulator and comparing the resulting output distributions against calibration targets.

Z-scores with pooled standard deviations were calculated to assess agreement between the calibrated model and population data,

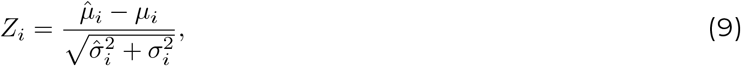

where 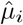 is the posterior mean and 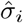 the posterior standard deviation. Outputs with |*Z*_*i*_ |≤1 are within one standard deviation of the data and thus considered in good agreement.

### 2.5 Surrogate model

The sensitivity and identifiability analyses and Hamiltonian Monte Carlo calibration all require a large amount of model evaluations, especially when exploring a large multidimensional parameter space as we do here. Despite the relatively low computational cost of our reduced-order model, this still poses a substantial computational burden. We therefore trained Gaussian Process Emulators (GPEs, one per output) as surrogate for our biophysics model. The GPEs were trained on the inputs and outputs of biophysics model evaluations, with the input space generated using Latin Hypercube Sampling of *N* = 4000 within the bounds of the prior distributions (Table 2). The AutoEmulate package^55^ was used to identify the optimal kernel and training configuration, and 5-fold cross validation and mean *R*^2^ across folds was used to identify the optimal configuration. This resulted in a matern 5/2 kernel with learning rate 0.2 and *R*^2^ values of 1.00 for all outputs except MAP (0.96) and *λ*_ED,LFW_ (0.98).

### 2.6 Assessing sex differences

HMC calibration yielded sex-specific posterior parameter sets, which we will refer to as the female model (*θ*_*F*_) and the male model (*θ*_*M*_). Whereas some studies have shown that most sex differences observed in the heart can be attributed to body size^56^, others suggest that the differences extend beyond size alone^30^. To determine whether sex differences in the calibrated parameters can be attributed to body size, two size-matched models were constructed by rescaling the body-size-dependent parameters of one sex to the body size of the other: *θ*_*F →M*_ denotes the female model at male body size, and *θ*_*M →F*_ the male model at female body size. The body-size-dependent parameters {*R*_as_, SBV, *A*_m,ref,*W*_, *W*_th,*W*_ }, for the five wall segments *W* ∈ { lfw, rfw, sw, la, ra }, were allometrically scaled as a function of body surface area (BSA)^57,58^,

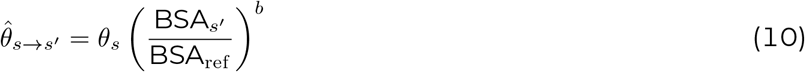

where *s* and *s*^*′*^ denote the two sexes, BSA_*F*_ =1.75 m^2^, BSA_*M*_ =2.03 m^2^, BSA_ref_ =1.9 m^2^ and *b* = −3*/*4 for *R*_as_ and *b* = 1 for SBV, *A*_m,ref_ and *W*_th_^57^. Since *θ*_*F →M*_ and *θ*_*M*_ are thereby expressed at the same body size, as are *θ*_*M→F*_ and *θ*_*F*_, differences remaining within either pair cannot be attributed to body size.

To assess statistical sex-differences, Welch’s t-test^59^, Benjamini-Hochberg false discovery rate (BH-FDR) correction^56,60^, and Cohen’s d analysis^56,61^ were conducted. Welch’s t-test assesses whether significant differences exist between the means of the independent groups without assuming equal variances. Raw p-values were corrected using BH-FDR to control the expected proportion of false discoveries among significant results. Cohen’s d quantifies the effect size by measuring the standardized difference between the means of two groups, independent of sample size. A value of 0 indicates equal means, and, using the Gignac & Szodorai thresholds^62^, d values were classified as small (0.2 ≤ |*d*| *<* 0.4), medium (0.4 ≤ |*d*| *<* 0.6) or large (|*d*| ≥ 0.6).

### 2.7 Code availability

All model code used to generate the results in this study is available on our lab’s GitHub page (http://www.github.com/BEATLabUCI). This repository includes: Python code of the reduced-order model; annotated Jupyter notebooks that provide guidance on using the code and reproducing all results and figures; CSV files containing all target data. We welcome anyone to submit issues via GitHub or e-mail to help us improve the code’s functionality and reproducibility.

## 3 RESULTS

### 3.1 Three-stage parameter pipeline for optimal model calibration

Our three-stage parameter reduction pipeline reduced the original 46-parameter space for the reduced-order model to 10 parameters in females and 9 parameters in males: removing 21 at the sensitivity stage, 14 at the FIM stage, and, 3 in females and 2 in males at the profile-likelihood stage (Table 2). Because identical priors and output choices (but not target values) were used for both sexes, the Sobol and FIM stages selected the same parameters for females and males. In contrast, the PL step is data-dependent; and it identified one unidentifiable parameter in males that is identifiable in females. The remaining parameters to be calibrated were not distributed evenly across model subsystems: cardiac geometry parameters *A*_*m*,ref,lfw_, *A*_*m*,ref,rfw_, *A*_*m*,ref,la_, *A*_*m*,ref,ra_; atrial active parameter *S*_f,act,a_, ventricular active parameters *S*_f,act,v_, *t*_ad,v_; and circulation parameters SBV, *r*_as_, and, *r*_lv_ for females.

Circulation, cardiac geometry, ventricular and atrial active contraction parameters were identified as the most influential during Sobol sensitivity analysis, whereas both ventricular and atrial passive parameters were found to be the least influential on the output variance (Fig. 2, A, B). Interestingly, ventricular outputs were more sensitive to the contractile parameters *t*_ad,v_ and *l*_s,ref,v_ than to *S*_f,act,v_, whereas atrial outputs were more sensitive to *S*_f,act,a_ and *l*_s,ref,a_ than to *t*_ad,a_.

**Figure 2:**
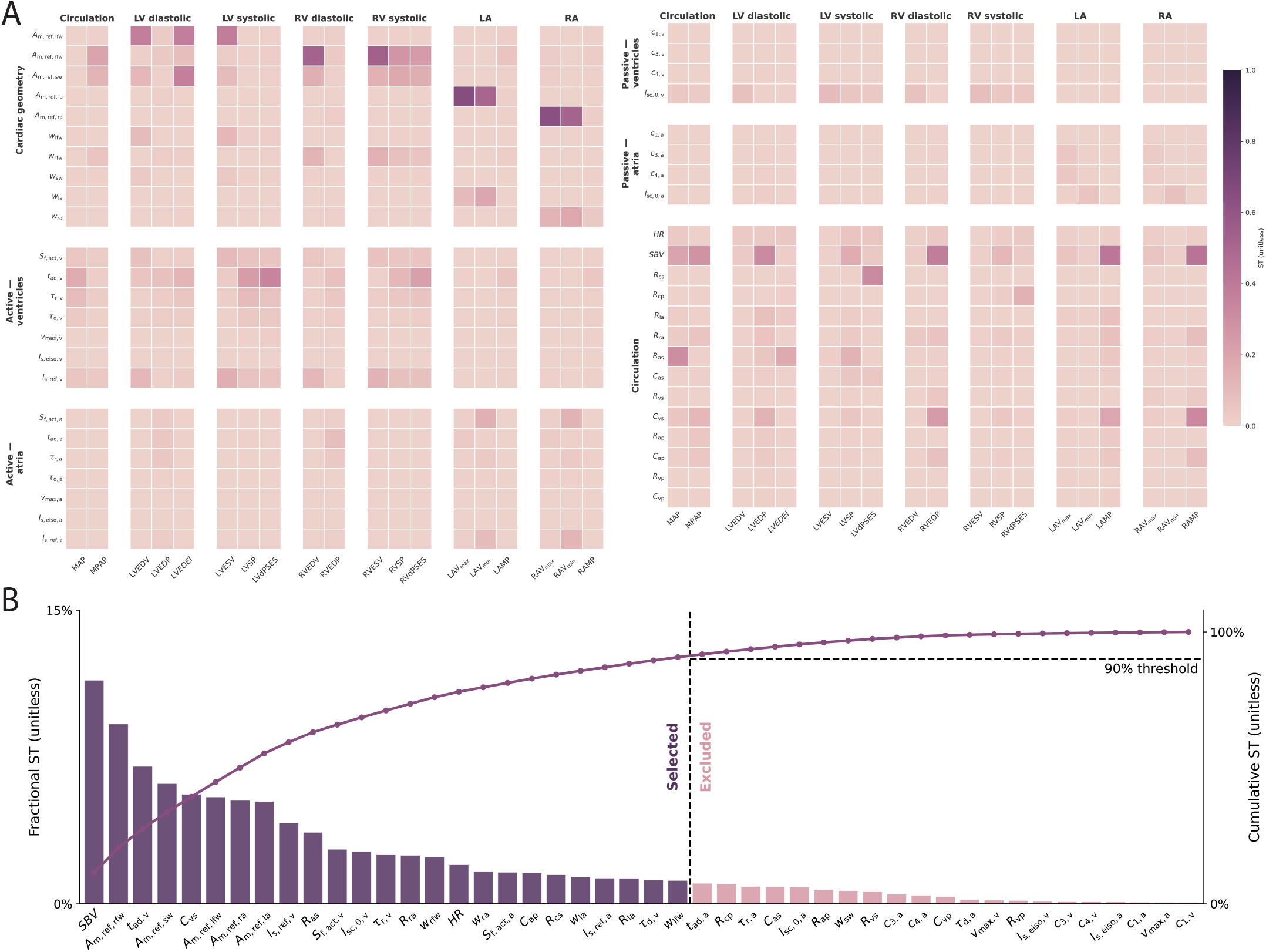
Sobol sensitivity was used to (A) compute total-order sensitivity indices across all model parameter and outputs, and (B) identify that 25 out of the 46 parameters accounted for 90% of the output variance.

FIM analysis identified the right free wall thickness as collinear with its corresponding wall area, with additional collinearity in circulation and ventricular active contraction parameters as shown in Fig. 3. The left and right free wall and left atrial wall thicknesses were found to be collinear; since the septal wall was already removed during SA, we then chose to also remove the right atrial wall thickness. The initial PL analysis (Fig. 4, A) revealed that the majority of the parameters were practically unidentifiable within the non-linear likelihood surface. PL after FIM-based collinearity removal (Fig. 4, B) showed that collinearity was the main driver of this unidentifiability, evidenced by tightened profiles and confidence intervals no longer hitting the lower and upper bounds. Despite their strong influence, circulation parameters remained among the most prone to collinearity or practical unidentifiability. This can be explained by the serial arrangement of the five-element RCRCR Windkessel models that describe the systemic and pulmonary circulations. Because the compartments are connected in series and the pressures and volumes included in the target data are too sparse to constrain every circulation parameter individually, individual resistances and compliances can trade off against one another without changing the observed outputs.

**Figure 3:**
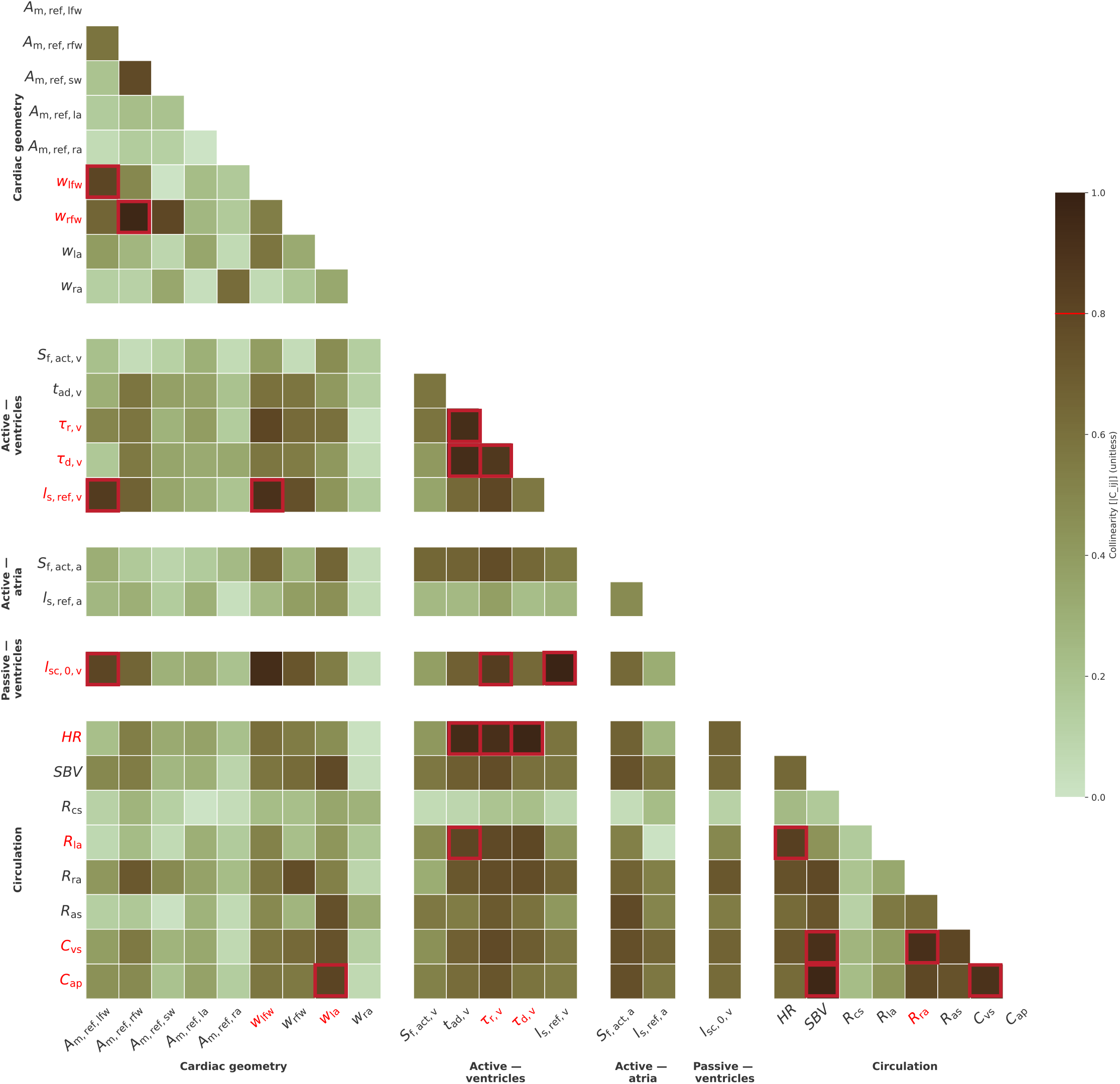
FIM analysis identified collinear pairs (*i, j*), defined as normalized correlation | *C*_*ij*_ | ≥0.8, indicated by red boxes. Of each correlated pair, the parameter with the lower first-order Sobol index *S*_1_ was excluded from calibration, indicated by a red label. Note that some red labels appear on both members of a pair, this occurs when a parameter is involved in multiple collinear interactions across different rows and columns. For example, HR is fixed due to collinearity with *t*_ad,v_ and *t*_r,v_, however, *t*_r,v_ is red-labelled due to its own collinearity with with *l*_sc,0,v_ and *t*_ad,v_. The heatmap represents the normalized cosine-similarity matrix, with the colorbar ranging from 0 to 1 indicating increasing collinearity.

**Figure 4:**
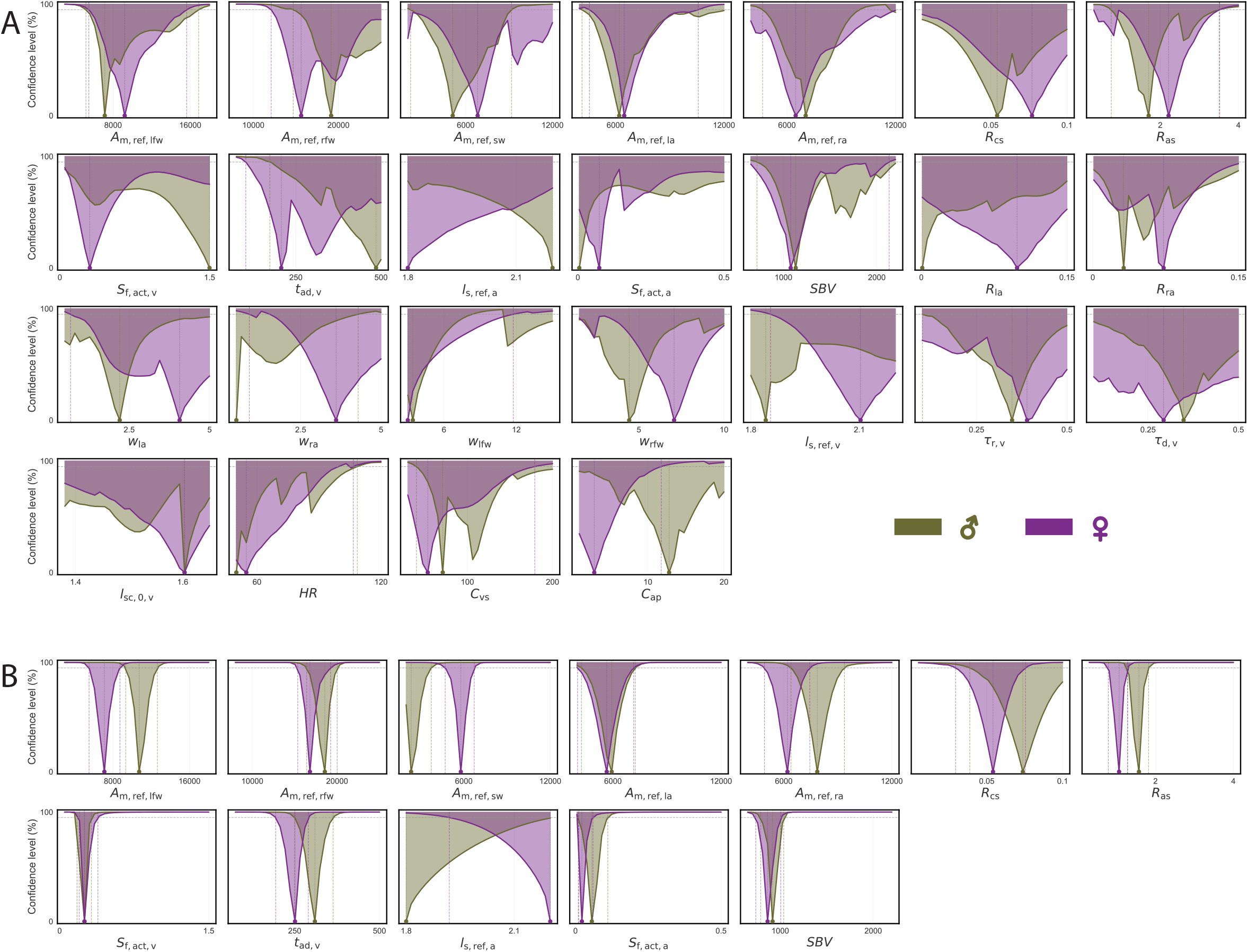
(A) PL analysis of all influential parameters indicate identifiability issues for most model parameters, indicating by the wide confidence intervals for both female (purple) and male (green) populations. (B) Repeating the analysis without the parameters identified as collinear by FIM, PL analysis indicates that *r*_lv_, *l*_s,ref,a_, and *A*_*m*,ref,sw_ remain unidentifiable for males and only *l*_s,ref,a_ remains unidentifiable for females

### 3.2 Calibrated parameter values differ between females and males

The parameters that were identified as sensitive and identifiable were calibrated for both sexes using Hamiltonian Monte Carlo. The pairwise relationships between the calibrated posterior parameters for females and males are visualized using kernel density plots (Figure 5). All calibrated parameters show less than 80% overlap between the sexes, indicating clear sex differences, especially for *S*_f,act,a_, *S*_f,act,v_ *A*_m,ref,ra_, *A*_m,ref,rfw_ and SBV, which each show less than 50% overlap. Additionally, the joint contours are predominantly oval-shaped, showing parameter independence. However, despite earlier efforts to address multicollinearity and identifiability in the pipeline, some collinearity persists: between *S*_f,act,v_ and other parameters. Finally, 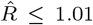 and ESS_bulk_ *>* 400 for all parameters.

**Figure 5:**
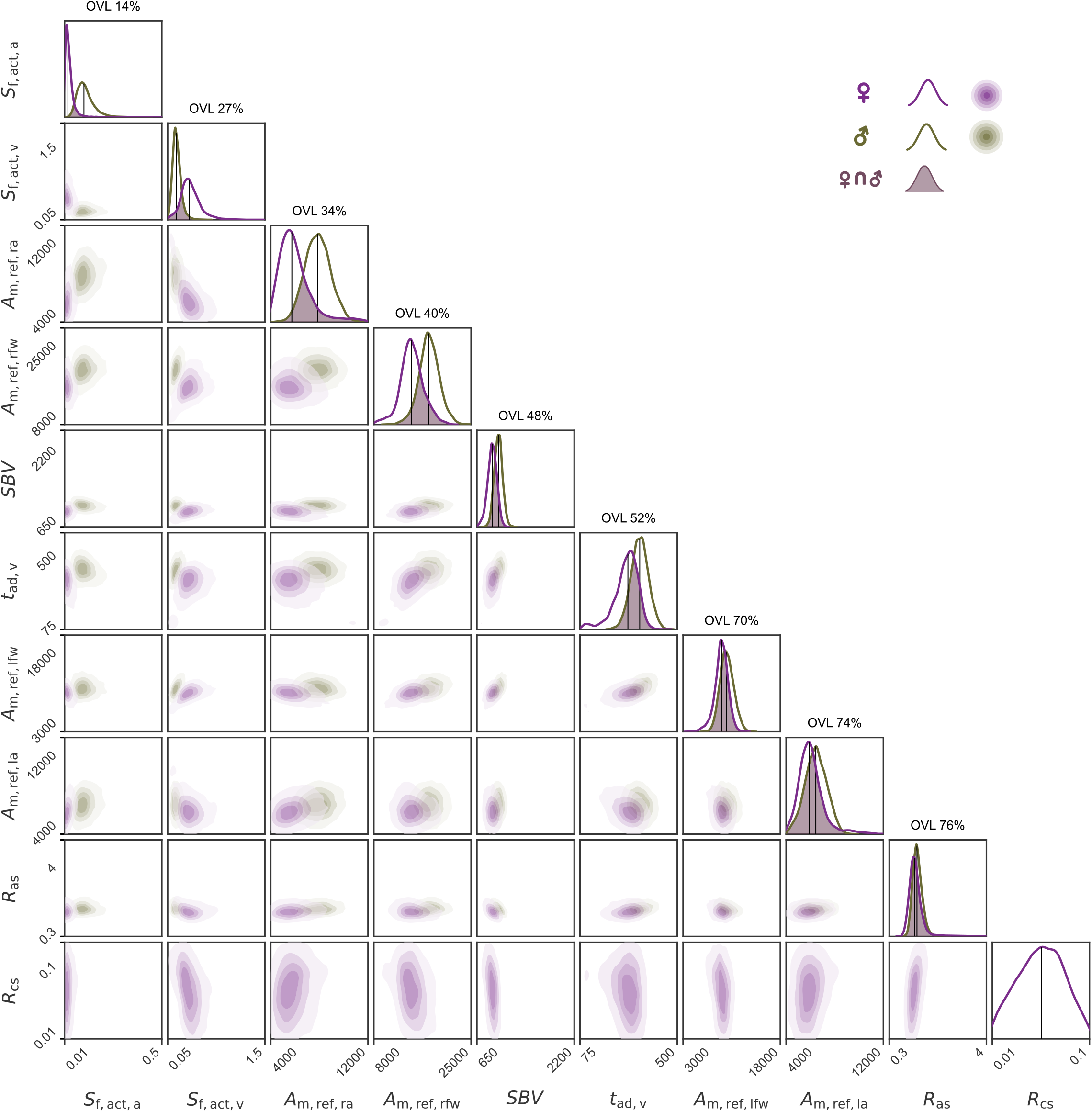
Kernel density estimates of the joint posterior distributions of the 10 calibrated parameters for female (purple) and male (green) populations. The off-diagonal panels show the pairwise posterior contours of each parameter pair, while the diagonal panels show the marginal posteriors for each sex along with the overlapping coefficient (OVL) value (arranged in increasing order of overlap), where greater values indicate higher overlap. The axis limits of each parameter are set to their prior range.

After calibration, the posterior distributions of the model outputs were compared against calibration targets to assess goodness of fit. For women, 18 out of the 19 target outputs were calibrated within |*Z* |≤1, while one target output (RAMP) had |*Z* | *>* 1. For men, 17 out of the 19 target outputs were calibrated within |*Z* | ≤1, while two target outputs (LVEDV and RVdPSES) had |*Z* |*>* 1. The output distributions (Figure 6) are unimodal, symmetric, and well-constrained with substantial overlap against the calibration targets across all outputs and both sexes.

**Figure 6:**
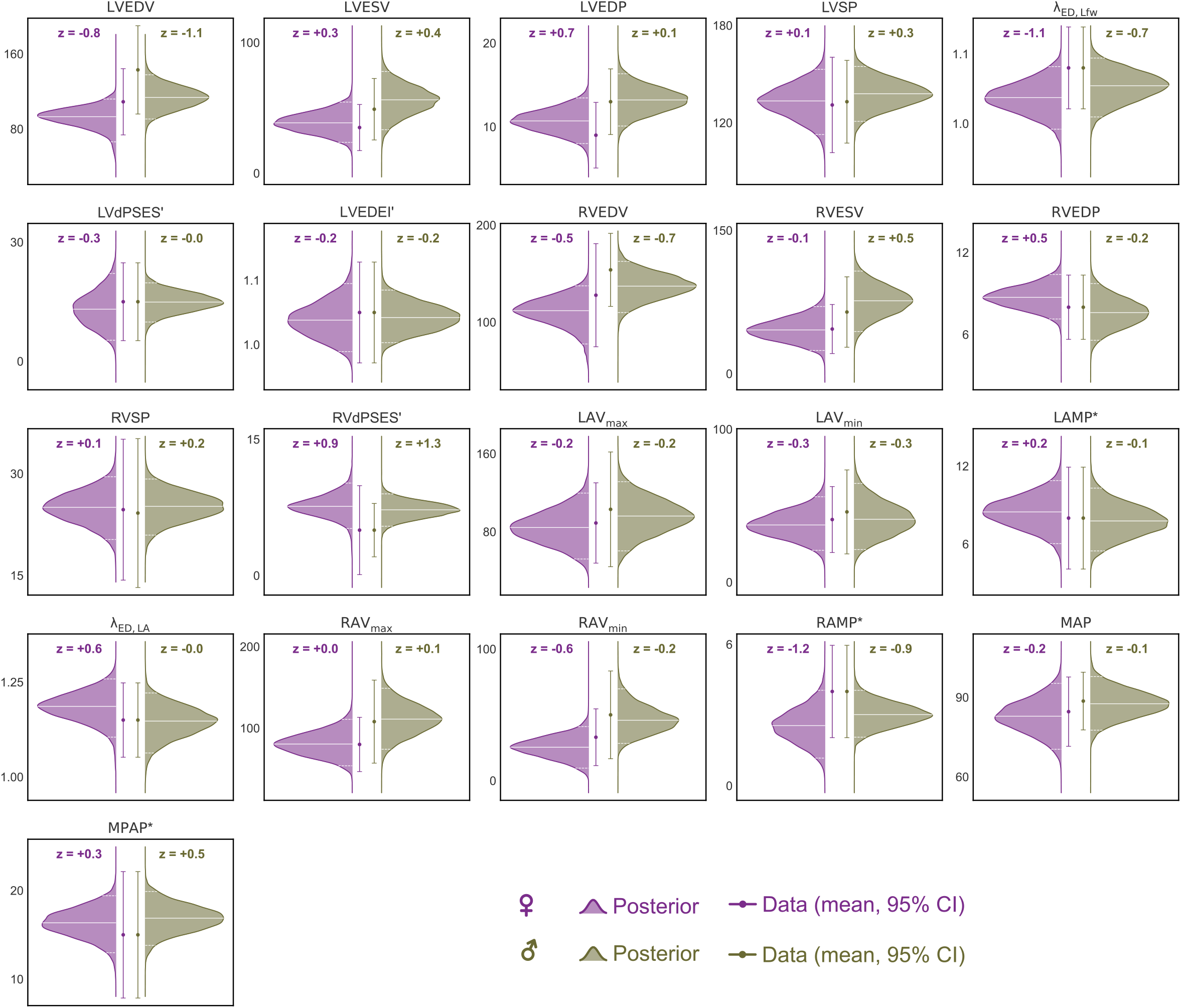
Posterior output distributions for female (purple) and male (green) populations across the 21 calibrated outputs. Data targets (mean and 95% CI) are indicated by whiskers, and each panel reports the z-score. Note that outputs marked with ^*′*^ show no sex-difference and those marked with *\** are not sex-specific.

### 3.3 Calibrated female and male simulations

We used 100 parameter sets from the posterior distribution to simulate PV loops for the 4 chambers across both sexes (Figure 7, A) show that the female heart operates at lower volumes, as indicated by the female curves being shifted leftward relative to the male curves in all chambers, with the shift being less pronounced in the LA. Interestingly, male ventricles generated more pressure, indicated by increased end-systolic pressure, despite a lower *S*_f,act,lv_ distribution. The Wiggers diagram Figure 7, B) further shows that the cardiac cycle is longer in males, potentially due to lower heart rates and greater ventricular activation duration *t*_ad,v_, a calibrated parameter. Additionally, the pressure difference between the left ventricle and systemic arteries during ejection was smaller in women than in men.

**Figure 7:**
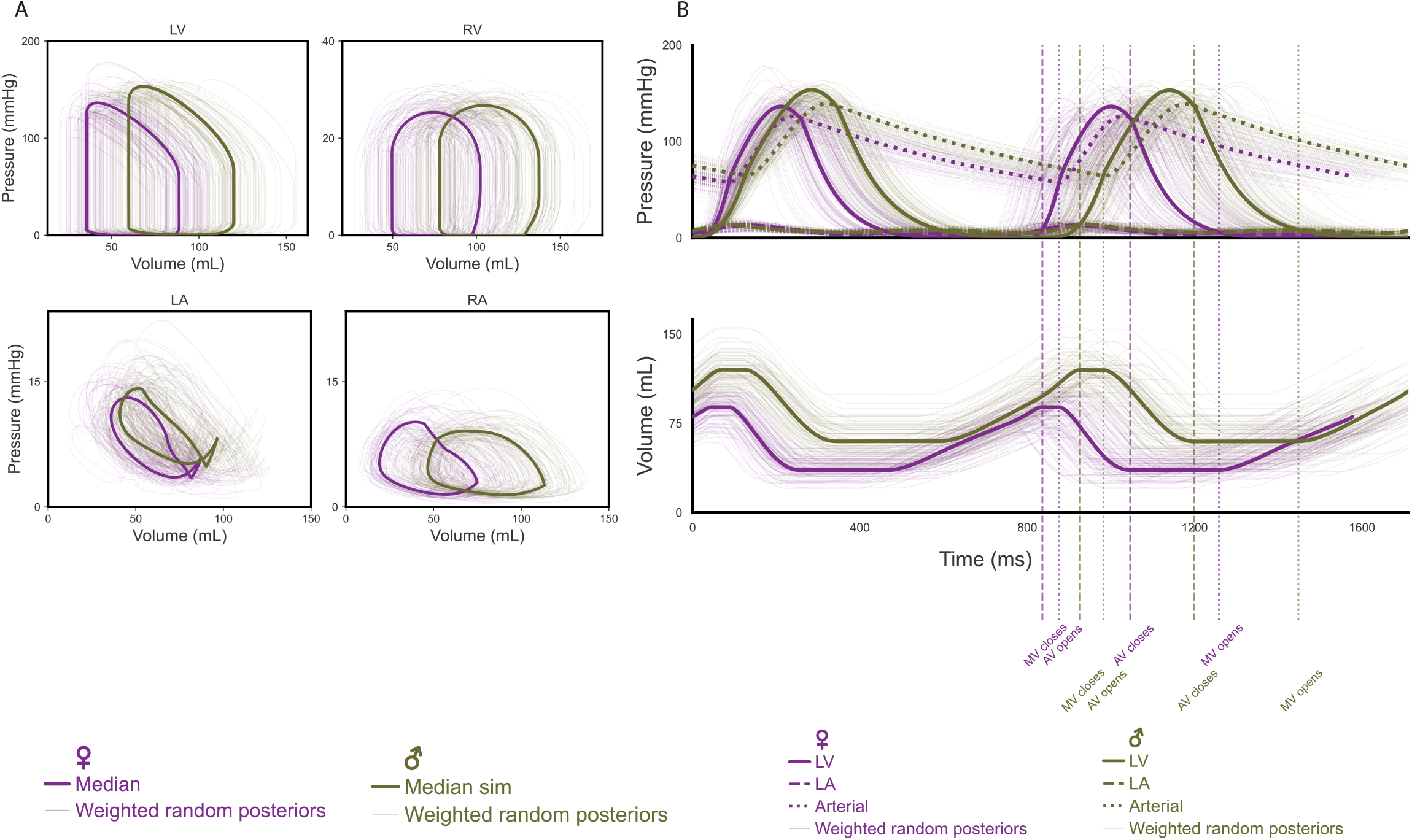
Hemodynamics were simulated using the calibrated female (purple) and male (green) posterior parameter distributions, showing (A) pressure-volume loops of the 4 heart chambers and, (B) The Wiggers diagram showing volumes, pressures and valve events across the cardiac cycle. In all panels, the median simulation is shown by the thicker line and the remaining simulations by the thinner lines.

### 3.4 Size does not explain all the sex differences

To assess whether body size explains differences between the posteriors, we compared *θ*_*F*_ and *θ*_*M*_ against allometrically-scaled *θ*_*M →F*_ and *θ*_*F→ M*_ (Section 2.6). Each model was used to simulate outputs that were held out from calibration (Figure 8). The sex differences separated into two groups. For cardiac output and PCWP, body-size correction substantially altered the difference: the male-female gap in LVCO and RVCO not only shrank but reversed direction under scaling, with females (*θ*_*F*_ and *θ*_*F→ M*_) reaching higher cardiac output—plausibly because their higher heart rate dominates once stroke volumes are equalized by scaling. The differences in PCWP likewise diminished with scaling, though PCWP remained higher in males (*θ*_*M*_ and *θ*_*M →F*_) throughout. In contrast, the differences in LVPER, LV and RV ejection fraction and maximum dP/dt were largely unaffected by scaling: females exhibited higher values in all cases, and both the direction and effect size largely persisted after allometric correction. The calibrated sex differences were consistent with reported values, with males showing 5% higher LVCO (versus 8% reported^63^) and females 7% higher EF (versus 5%^64^). Because effect sizes remained non-zero across all scaled comparisons, body size accounts for some, but not all, of the observed sex differences.

**Figure 8:**
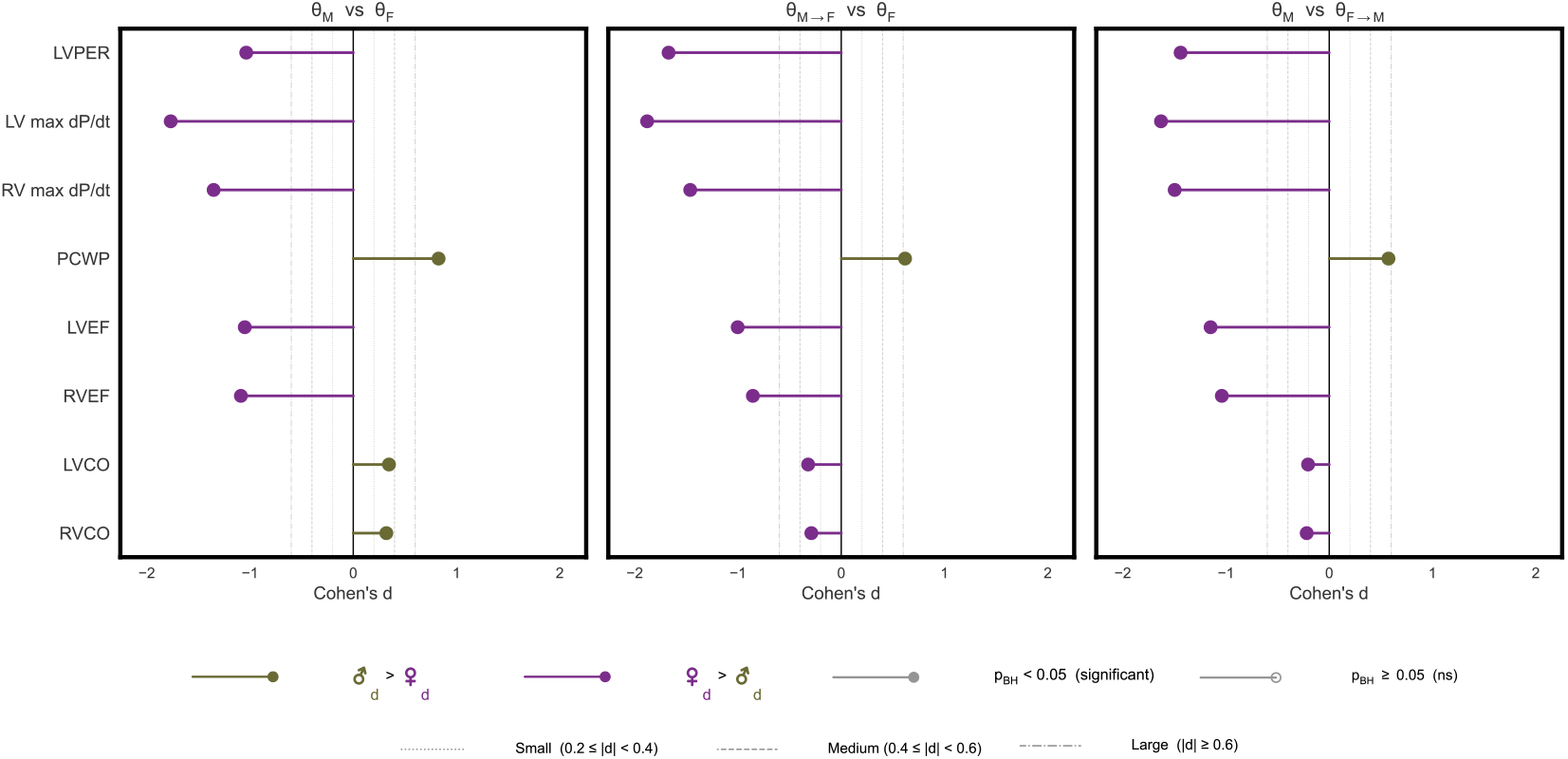
Assessing statistical sex differences using Cohen’s d analysis on outputs held out from calibration between female (*θ*_*F*_), male (*θ*_*M*_), size-matched female (*θ*_*M→F*_) and size-matched male (*θ*_*F→ M*_). Differences are directionally computed as male minus female, or *θ*_*M→F*_ minus F, or M minus *θ*_*F→ M*_, with positive values indicating a higher value in the first term of each comparison. LVPER, LV Peak Ejection Rate; LV max dP/dt, Maximum Rate of Pressure Rise in the LV; RV max dP/dt, Maximum Rate of Pressure Rise in the RV; PCWP, Pulmonary Capillary Wedge Pressure; LVEF, LV Ejection Fraction; RVEF, RV Ejection Fraction; LVCO, LV Cardiac Output; RVCO, RV Cardiac Output.

## 4 DISCUSSION

Reduced-order cardiovascular models retain essential physiological behavior at substantially lower computational cost than high-fidelity counterparts, and are increasingly used in academia as well as the medical device and pharmaceutical industries for both mechanistic and patient-specific studies. Sensitivity analysis, identifiability screening, and Bayesian inference have each been applied to such models individually, but they are rarely combined, and where they are, the aim is typically to personalize a model to an individual patient or disease state rather than to characterize healthy physiology. In this work, we combined all three to calibrate a reduced-order model for healthy female and male populations, establishing sex-specific posterior parameter distributions and quantifying how much of the observed sex differences can be explained by body size. The resulting parameter distributions provide reference ranges for future mechanistic and patient-specific simulations.

### 4.1 Sex differences in calibrated parameters exceed body size

All calibrated parameter distributions show <80% overlap between women and men (Figure 5), highlighting the importance of sex-specific parameterization. Among the parameter posteriors with low overlap *S*_f,act_, *A*_m,ref,rfw_ and SBV - the latter two were also among the most influential (Figure 2) and are expected to be related to body size. However, after allometric scaling, significant effect sizes persisted, suggesting that body size alone does not fully account for the observed sex differences (Figure 8). One possible explanation is heart rate, a latent parameter with sex-specific values used during calibration that was not allometrically scaled in this study and is explicitly used to calculate outputs such as LVCO and RVCO, and implicitly impacts PCWP. Scaling already reduced these effect sizes, so these differences might diminish further if heart rate was also scaled, along with LVPER, EF and max dP/dt which are currently scaling agnostic.

### 4.2 A three-stage pipeline improves identifiability before calibration

The three stage parameter subset selection pipeline reduced the number of parameters from 46 to 10 in females and 9 in males. Each stage removed parameters for a distinct reason: low output sensitivity (46 *→*25 parameters, Fig. 2), collinearity with other parameters (25 *→*14 parameters, Fig. 3), and structural or practical unidentifiability persisting in the absence of collinearity (14*→* 9-10 parameters, Fig. 4); no single analysis by itself would therefore have sufficed. The first two stages are also computationally inexpensive: since the FIM (Eq. 3) is assembled from the model evaluations already required for the sensitivity analysis, collinearity screening adds no further model evaluations, and the two stages together account for 35 of the 36 (female) and 37 (male), removed parameters in both sexes. Profile likelihood identified 2 parameters in women and 3 parameters in men to be removed, in this case because of practical unidentifiability, but it can also detect structurally unidentifiable parameters that could be present in a different model, parameter, and/or output combinations. Of note here *A*_*m*,ref,sw_ is unidentifiable in males during PL, and is fixed as a ratio of *A*_*m*,ref,lfw_. This was also done in females both for geometric calibration consistency and as we expect a larger value for males rather than females which is not seen in the PL (Fig. 4, B). In males *r*_lv_ was found to be unidentifiable, with the results suggesting that a higher prior may have resolved the issue. However, as the prior range is already quite generous, adjusting this value would require restarting the calibration pipeline from the sensitivity analysis stage. Using this pipeline, we obtained unimodal and well-constrained joint posteriors for both female and male populations (Fig. 5). The *S*_f,act,v_ profile seems to be similar in PL for women and men Fig. 4, B), however, this parameter also has one of the least overlap between the sexes in their posterior (Fig. 5). This could be due to PL profiling one parameter at a time and missing the joint influence of other parameters.

The rigorous emphasis on parameter identifiability distinguishes our methodology from much of the existing literature. Many deterministic^17–20^ and Bayesian^13,16,24,26–29,65–68^ calibration studies exist for high fidelity and reduced-order cardiovascular modeling alike. Many of these studies include sensitivity analysis without identifiability analyses^13,16,27,56,66,69^, leaving potential parameter redundancy unaddressed prior to calibration. Unaddressed parameter unidentifiability manifests differently across the two calibration frameworks: deterministic calibrations could yield non-unique solutions where various parameter combinations fit the data equally well; Bayesian calibrations could yield broad, strongly correlated joint posteriors that are difficult to interpret and can impede sampler convergence.

PL alone is employed in a handful of studies in biology^50^ and cardiovascular model calibration^17,18,68,70–72^, but fewer still combine both FIM and PL^24,28,29,71,72^. Identifiability analyses incorporating both FIM and PL are particularly valuable: while PL by itself can reveal unidentifiability issues (Figure 4, A), combining it with FIM to remove collinear pairs driving practical unidentifiability provides a better-defined region for likelihood estimation, resulting in tighter confidence intervals (Figure 4, B). Schiavazzi et al.^29^ represent the closest methodological predecessor to the present work, employing SA-FIM-PL pre-calibration and Bayesian MCMC to calibrate a 0-D circulation model to diseased single-ventricle data without sex-stratification.

Finally, the reported parameter selection and their posterior distributions depend on our particular combination of model and data, e.g. the serial RCRCR model employed here requires local pressure measurements to identify all resistances and capacitances. The calibration pipeline, however, is model- and data-agnostic and can be followed by others to obtain well-informed parameter distributions for their own models. Therefore, this study does not only offer sex-specific reference parameters for reduced-order cardiovascular models, but also a transferable procedure for robust parameter identification.

### 4.3 Population-level calibration from sex-balanced healthy cohorts

Comprehensive, sex-stratified, four-chamber measurements from a single healthy cohort are scarce, and calibration data were therefore aggregated across 11 healthy cohorts. Of these, 10 comprise between 12 and 850 participants, depending on the measurement, with a well-balanced female representation of 40-60%. The remaining three cohorts include no female participants, a limitation addressed in Section 4.4. Sex-specific parameterization of reduced-order cardiovascular models is lacking. Existing reduced-order modeling studies that include female data do not sex-stratify their parameter estimates^17–19,29^, and of these, only one used healthy human female data^17^.

This gap persists in high-fidelity modeling studies. Rodero et al.^65,69^ parameterized a high-fidelity model using 19 healthy participants (30% female), generating 1,000 synthetic cases; however, the synthetic cohort is not stratified by sex. Solis-Lemus et al.^56^ represents a significant step forward as the first study employing a sex-balanced cohort with a four-chamber electromechanical high-fidelity model in both healthy and diseased participants, but it did not include sensitivity or identifiability analyses.

Since we combined datasets to obtain more measurements than are commonly available, the sex-specific posterior distributions in Figure 5 and Table 2 can serve as reference values for parameterizing reduced-order model studies, especially when parameters are not identifiable from the available data. For example, invasive pressure measurements are often unavailable in routine clinical data, so circulatory parameters distributions identified in this study can be used as informed priors or latent parameters in patient-specific calibrations. More broadly, our pipeline can be re-run to parameterize independent studies, as the Windkessel-based RCRCR circulation model employed here is widely used^9,13,16,34,73–76^ and the Triseg formulation is arguably the most common reduced-order model of the heart^10^.

### 4.4 Limitations

This study has limitations at both the data and model levels, the most notable of which are discussed below.

At the data level, our calibration relies on healthy subject data drawn from pooled studies, which introduces several limitations. First, the binary sex categorization used does not account for intersex, non-binary, or transgender identities, reflecting broader societal limitations in the available data. Second, there are inherent differences across studies: data were collected using different modalities (such as MRI and ultrasound), span a wide age range (18-90 years), and were collected in different regions (USA, Canada, UK, and Italy). The diversity of the pooled data, however, can be viewed as a strength, lending the calibrated reference ranges for women and men a degree of generalizability that a single-cohort study could not provide. Third, some measurements, such as atrial wall thickness and some pressures, are scarce due to the invasive nature of data collection or their typical focus on disease states, resulting in low or unknown sample sizes. Lacking sex-stratified measurements, we therefore relied on male studies for the left^36^ and right atrium^42,46^ and RVEDP^41,42^. Note that Chemla et al.^41^ suggest that there are no sex differences in RVEDP. Fourth, a further consequence of using pooled cohort-level statistics is that the posterior distribution widths shown in Figure 5 conflate biological variability with measurement error and inter-study heterogeneity. Consequently, the reported overlap coefficients between sex-specific posteriors reflect a composite variance and may not correspond directly to the biological separation of the two populations.

At the model level, several limitations remain. First, every model rests on simplifications of reality, and ours is no exception: like any reduced-order model, ours captures cardiovascular physiology through a deliberately simplified representation. Therefore, several physiological mechanisms that may feature sex differences are not considered, including autonomic and baroreflex regulation^77^ and heart shape^78^. Second, the output likelihood (Eq. 8) treats outputs as independent, neglecting any potential correlations across outputs^68^. Because our targets are pooled marginal statistics, the covariances required for a multivariate likelihood were unavailable. This also motivated the choice of a separate GPE per output instead of a multiprocess GPE, which increased the surrogate model robustness at the cost of neglecting output covariance. Third, some model parameters remained practically unidentifiable even after eliminating non-sensitivity and collinear parameters (Fig. 4B). While practical unidentifiability could be resolved by adding more calibration targets, it could here also reflect a limitation of the reduced-order model. For example, Fig. 5 shows that a wide range of septal wall configurations and either left or right free wall configurations can produce similar outcomes, even after calibrating to the LV Eccentricity index in an attempt to identify the optimal Triseg configuration.

### 4.5 Conclusions

In this study we present a framework integrating a rigorous three-stage pre-calibration pipeline—global sensitivity analysis using Sobol’s method, FIM-based collinearity analysis, and profile likelihood identifiability analysis with parameter re-optimisation—followed by Bayesian inference to estimate sex-specific cardiovascular parameter distributions from a reduced-order model trained on a sex-balanced healthy cohort. To our knowledge, this is the first reduced-order cardiovascular model to combine these elements, addressing the dual gap of sex imbalance and insufficient pre-calibration rigor in the existing literature. Our results demonstrate that sex differences in calibrated parameters exist and extend beyond body size alone. These results establish healthy sex-specific reference parameter distributions and contribute to more inclusive cardiovascular computational modeling.

## ACKNOWLEDGEMENTS

This work was supported by the American Heart Association Career Development Award (https://doi.org/10.58275/AHA.25CDA1452893.pc.gr.229647) and the Society of Hellman Fellows, awarded to PJAO, which also provided partial support for AL. The funders had no role in study design, data collection and analysis, decision to publish, or preparation of the manuscript. We thank our lab members for their helpful discussions and feedback.

## S1 MODEL DESCRIPTION AND IMPLEMENTATION

This supplement describes the reduced-order model and its implementation in detail. Our model is not new; its components are built from foundational work by many others, and comparable versions were published by authors from this manuscript^16,34,79^. To avoid confusion that could arise due to updates in the model between papers, this document describes the specific model formulation used in this manuscript, and the specific Python code used in this manuscript *will be made publicly available on GitHub upon publication*.

Note that the model code repository includes some features that are not used in this manuscript. First, the model implementation supports multiple wall patches per wall segment, following the MultiPatch formulation of Walmsley *et al*. (2015)^11^. Since all simulations presented in this study use a single patch per wall segment, the patch level is omitted from the description below and all quantities are defined per wall segment *W* . For an implementation with multiple patches per wall segment, the reader is referred to Walmsley *et al*.^11^ for details. Second, the code repository includes capabilities to simulate mechanics-driven cardiac growth. Readers interested using our model code for this purpose are referred to Jones *et al*. for volume overload^79^ and Oomen *et al*. for left bundle branch block and cardiac resynchronization therapy^34^.

### S1.1 Volumes and pressures

The hemodynamic circulation (Fig. 1) is modeled by a lumped-parameter circuit model, previously described by^9^. The lumped-parameter circulation comprises four vascular compartments *V* ∈ { as, vs, ap, vp } and four cardiac cavities *C* ∈ { la, lv, ra, rv } (Figure 1). The stressed blood volume SBV is defined as the total volume that raises pressure above zero in the vascular compartments, together with a lblood contained in the cardiac cavities (Eq. S1):

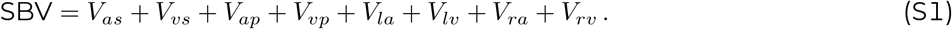

At the start of the simulation, the stressed blood volume is distributed across compartments according to an initial guess **k** = [*k*_*q*_]_*q∈ V∪C*_ with Σ_*q*_ *k*_*q*_ = 1, such that the initial volume in each compartment is given by Eq. S2:

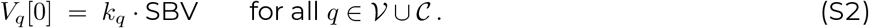

When the model is run for successive cardiac cycles at a constant heart rate, the volume distribution converges to a periodic steady state in which the change in distribution between consecutive cycles becomes negligible. A good initial guess closely approximates this converged distribution, thereby reducing the number of simulated cardiac cycles before the haemodynamic simulation can begin.

The compartment pressures *P*_*q*_ are computed differently for the vasculature and cardiac cavities. For the vascular compartments, pressure is given by the ratio of current blood volume *V*_*q*_ to the compartment capacitance *C*_*q*_ (Eq. S3):

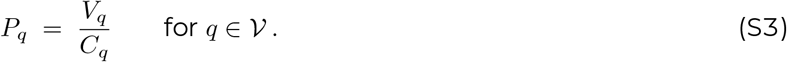

For the cardiac cavities, pressure is derived from Laplace’s law for a thin-walled sphere (Eq. S4), relating the midwall tension *T*_*m,W*_ and midwall radius *r*_*m,W*_ of the corresponding wall segment (described in Section 2.2):

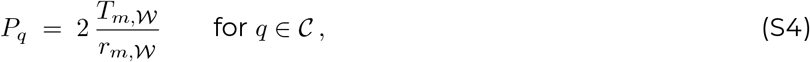

where the mapping between cavity *q* and wall segment *W* is lv→LFW, rv→RFW, la→LA, ra→RA.

Blood flows between adjacent compartments along the pressure gradient, inversely proportional to the resistance between them. For a compartment *q* receiving flow from an upstream compartment *u* across resistance *R*_*u*_ and delivering flow to a downstream compartment *d* across resistance *R*_*q*_, the volume is updated according to Eq. S5:

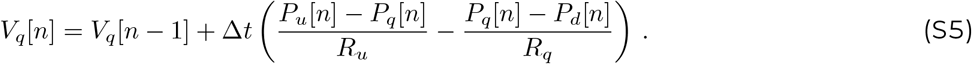

This general form applies to all vascular compartments, with upstream–downstream connectivity and corresponding resistances following the circuit shown in Figure 1. For the atria, ventricles and arteries, the same structure holds, but a unidirectional flow condition is imposed at each heart valve using the Iverson bracket [·], which evaluates to 1 when the enclosed condition is true and to 0 otherwise. For example, the left ventricular volume update is given by Eq. S6:

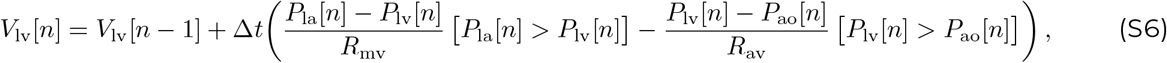

where *R*_mv_ and *R*_av_ denote the mitral and aortic valve resistances, respectively.

### S1.2 Sarcomere contraction model

Throughout the following, parameters that may differ between cardiac wall segments carry the subscript *W*.

#### S1.2.1 Contractile element length

The mechanics of the wall segments are described by a Hill-type sarcomere contraction model first described by Arts et al.^80^, later defined for the TriSeg model by Lumens et al.^81^. The sarcomere is modeled as a passive element of length *l*_s_ in parallel with a contractile element (length *l*_sc_) and an elastic element (length *l*_se_) arranged in series, such that *l*_s_ = *l*_se_ + *l*_sc_. In what follows, the subscript *X* ∈ { *a, v* } denotes the wall type (atrial or ventricular), and all chamber-specific parameters carry this subscript.

The total sarcomere length is determined by the myofibre stretch *λ*_f,*W*_ (derived from wall geometry, Section S1.4) and the reference sarcomere length at zero strain *l*_s,ref, *X*_ . The elastic element length then follows from the series relationship (Eq. S7):

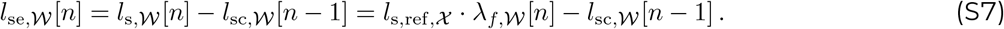

The contractile element length is updated at each time step as a function of the elastic element length relative to its length during isometric contraction *l*_se,iso,*X*_ (Eq. S8):

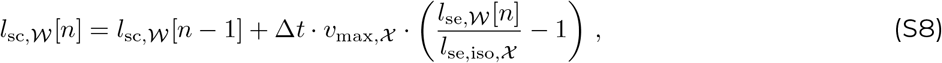

where *v*_max,*X*_ is the unloaded sarcomere shortening velocity. During isometric contraction (*l*_se,*W*_ = *l*_se,iso, *X*_) the contractile element length remains unchanged; it shortens when external load stretches the elastic element beyond this reference and lengthens when the elastic element is compressed below it.

#### S1.2.2 Contractility

The contractility *c*_*W*_ is a lumped variable representing the intracellular calcium transient that drives active force generation. Its evolution is governed by an excitation term *c*^+^ and a relaxation term *c*^*−*^ (Eq. S9):

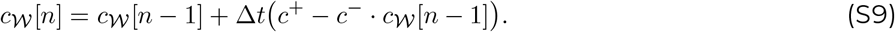

The excitation term *c*^+^ is the product of two factors: a rise function *f*_rise_ that models the calcium transient following electrical stimulation, and a length factor *f*_l_ that modulates the excitation relative to the contractile element length, scaled by the rise time constant *t*_r, *X*_ and the activation duration *T*_ad, *X*_ (Eq. S10):

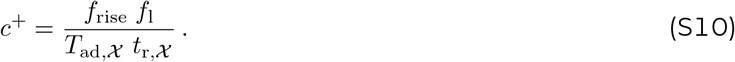

The rise function accumulates contributions from recent electrical activation events. Given a sequence of *K* activation times *t*_act,*k*_, only those within the window *t* − 8 *T*_ad,*X*_ *t*_r,*X*_ *< t*_act,*k*_ *< t* contribute (Eq. S11):

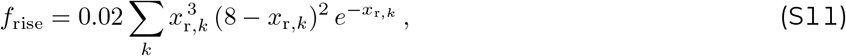

where the normalised time elapsed since activation *k* is (Eq. S12):

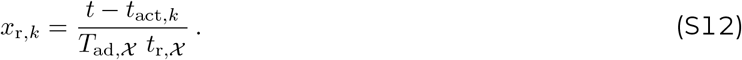

The length factor *f*_l_ scales the excitation as a function of the contractile element length relative to its resting length *l*_sc,0,*X*_ (Eq. S13):

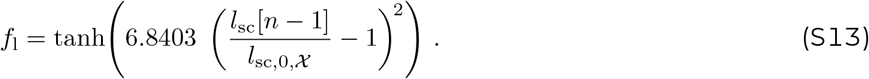

The relaxation term *c*^*−*^ is gated by a decay function *g*_decay_ that determines when relaxation begins (Eq. S14):

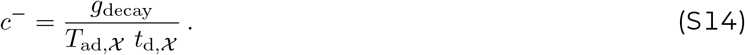

The decay function transitions smoothly from zero to one through a clipped sinusoidal ramp (Eq. S15):

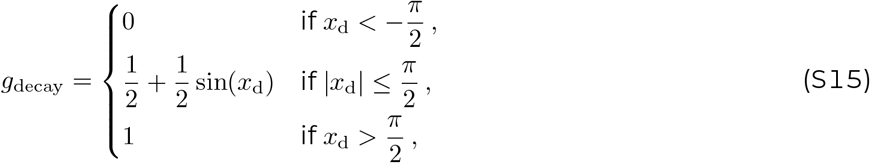

where the decay argument *x*_d_ offsets the onset of relaxation by a length-dependent term, so that longer sarcomeres sustain activation for a longer period (Eq. S16):

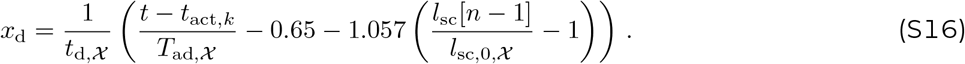

The decay term is computed with the most recent activation time *t*_act,*k*_ ≤ *t < t*_act,*k*+1_. While *c*^+^ is active only briefly after each electrical stimulus, *c*^*−*^ acts continuously once *g*_decay_ opens, drawing contractility back toward zero until the next activation event.

### S1.3 Atrial myofiber mechanics

Throughout this section, *W* denotes an atrial wall segment, *W* ∈ { LA, RA}, and *C* an atrial cavity enclosed by that wall segment, *C* ∈ { la, ra} . The atrial myofiber mechanics follow from the stretch required for the atrial wall to enclose its cavity volume. Since each atrium is modeled as a separate sphere, its geometry is determined by its cavity volume alone, and no equilibrium of tension between wall segments is required, unlike in the TriSeg formulation used for the ventricles (Section S1.4).

Under the thin-wall approximation, the wall volume of a spherical wall segment is the product of its reference midwall area *A*_*m*,ref,*W*_ (Table 2) and its wall thickness *W*_th,*W*_ (Table 2), *V*_*w,W*_ = *A*_*m*,ref,*W*_ · *W*_th,*W*_, and the midwall encloses the cavity and half of the wall volume, 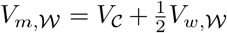 . Since both *A*_*m*,ref,*W*_ and *W*_th,*W*_ are constant, the midwall radius follows at each time step from the atrial cavity volume alone,

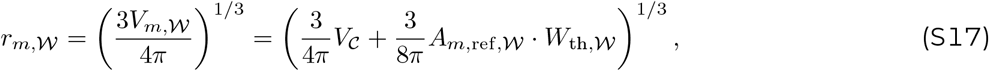

and, from the geometry of a sphere, the midwall area follows from the midwall radius as 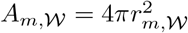 . The myofiber stretch was obtained as the square root of the ratio of the current midwall area to the reference midwall area:

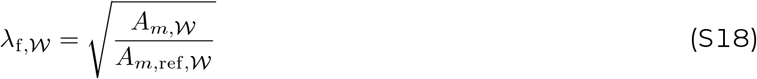

The isometric active stress was obtained from the contractility *c*_*W*_ (Eq. S9), the contractile element length *l*_sc,*W*_ (Eq. S8), the zero-stress atrial sarcomere length *l*_sc,0,*a*_ (Table 2) and the peak atrial active stress *S*_f,act,a_ (Table 2):

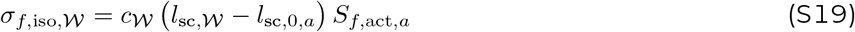

The total myofiber stress *σ*_*f,W*_ was modeled as the sum of an active and a passive stress component. The active stress was proportional to the extension of the series elastic sarcomere, *λ*_*f, W*_ · *l*_s,ref,*a*_ − *l*_sc, *W*_, with the atrial sarcomere reference length *l*_s,ref,*a*_ (Table 2), normalised by the atrial sarcomere shortening at isovolumetric contraction *l*_se,iso,*a*_ (Table 2). The passive stress represented the elastic response of the myocardial tissue to deformation and was determined by the atrial passive stiffness constants *c*_1,*a*_, *c*_3,*a*_, and *c*_4,*a*_ (Table 2)^34^:

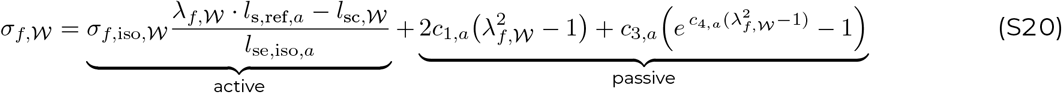

The atrial midwall tension followed from the myofiber stress and the wall geometry:

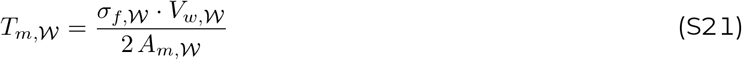

The output is an updated midwall tension *T*_*m, W*_ and midwall radius *r*_*m, W*_ of the wall segment *W* for the cardiac cavity pressure computation (Eq. S4), and an updated myofiber stretch *λ*_f, *W*_ for the contractile element length computation (Eq. S8).

### S1.4 Ventricular myofiber mechanics

Throughout this section, *W* denotes a ventricular wall segment, *W* ∈ { LFW, RFW, SW} . The geometry of each atrium follows from its cavity volume alone (Section S1.3). For the ventricles, described using the TriSeg formulation, this is not the case. Three wall segments enclose the left and right ventricular cavity volumes and share the septal wall, so that the position of a single segment affects the midwall area, and thereby the midwall tension, of all three. At each time step, the ventricular geometry is therefore found by adjusting the curvature of the three spherical wall segments until both cavity volumes are enclosed and the midwall tensions cancel at the junction circle along which the segments meet.

This search requires the midwall tension of each wall segment for every candidate geometry, but the relation between midwall area and midwall tension is nonlinear and costly to evaluate. Since the simulation time step is only a small fraction of the cardiac cycle, the midwall area of each wall segment changes only slightly between time step, and we assume the area–tension relation to be linear over this interval. The ventricular myofiber mechanics were therefore updated using a predictor-corrector scheme.

In the predictor step, the midwall area and midwall tension of each wall segment are obtained from its contractile element length alone, and are in general not compatible with the chamber volumes at the current time step. With this prediction as the expansion point, the area–tension relation was approximated by a first-order Taylor expansion, so that each wall segment is represented by its predicted midwall area and midwall tension together with the local slope of the relation. In the corrector step, the ventricular geometry satisfying both the cavity volumes and the equilibrium at the junction circle is determined with a Newton–Raphson solver, initialised at the geometry of the previous time step. Were the midwall tension evaluated directly, each iteration of the solver would require the nonlinear area–tension relation to be recomputed for all three wall segments. Since the tensions instead follow from the Taylor expansion, the relation is evaluated once per time step, in the predictor step, rather than at every iteration. The myofiber mechanics are then recovered at the converged geometry.

The scheme takes as input the contractile element length of each wall segment, the chamber volumes, and the ventricular geometry of the previous time step, and returns the midwall tension *T*_*m,W*_ and midwall radius *r*_*m,W*_ of the wall segment W for the cardiac cavity pressure computation (Eq. S4), together with the myofiber stretch *λ*_*f,W*_ for the contractile element length computation (Eq. S8).

#### S1.4.1 Predictor

For each wall segment, the predicted myofiber stretch 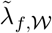 was obtained from the contractile element length *l*_sc,*W*_ (Eq. S8), normalised by the ventricular sarcomere reference length *l*_s,ref,*v*_ (Table 2):

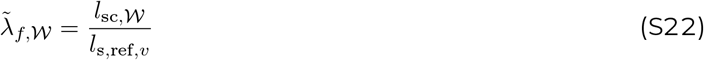

The isometric active stress was obtained from the contractility *c*_*W*_ (Eq. S9), contractile element length *l*_sc,*W*_ (Eq. S7), the ventricular zero-stress sarcomere length *l*_sc,0,*v*_ (Table 2) and the ventricular peak active stress *S*_f,act,v_ (Table 2):

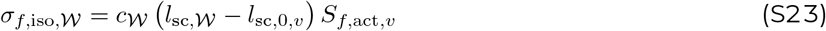

Predicted total myofiber stress 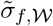 was modeled as a mixture of active and passive stress. The active stress was proportional to the extension of the series elastic sarcomere, 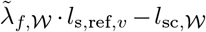, with the ventricular sarcomere reference length *l*_s,ref,*v*_ (Table 2), normalised by the ventricular sarcomere shortening at isovolumetric contraction *l*_se,iso,*v*_ (Table 2). The passive stress represented the elastic response of the myocardial tissue to deformation and was determined by the ventricular passive stiffness constants *c*_1,*v*_, *c*_3,*v*_, and *c*_4,*v*_ (Table 2):

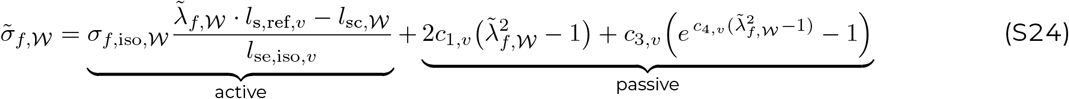

From the predicted stretch, the predicted midwall area *Ã*_*m,W*_ was obtained using the reference midwall area *A*_*m*,ref,*W*_ (Table 2):

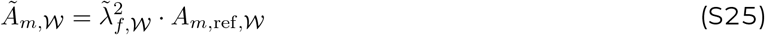

Under the thin-wall approximation, the wall volume of a spherical wall segment is the product of the reference midwall area *A*_*m*,ref,*W*_ (Table 2) and the wall thickness *W*_th,*W*_ (Table 2), *V*_*w,W*_ = *A*_*m*,ref,*W*_ · *W*_th,*W*_ . The predicted midwall tension 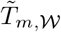 followed from the myofiber stress and the wall geometry:

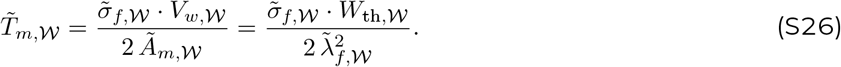

#### S1.4.2 Corrector

##### Linearized area-tension relation

The midwall tension was approximated by its first-order Taylor expansion about the predicted state,

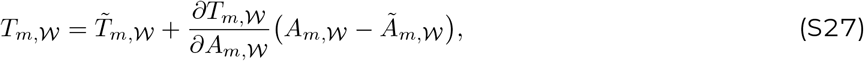

Since the midwall tension follows from the myofiber stress and the midwall area as *T*_*m, W*_ = *σ*_*f, W*_ · *V*_*w, W*_ */*(2 *A*_*m, W*_), and the myofiber stretch from the midwall area as 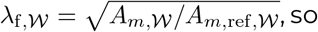, the slope follows by differentiation as

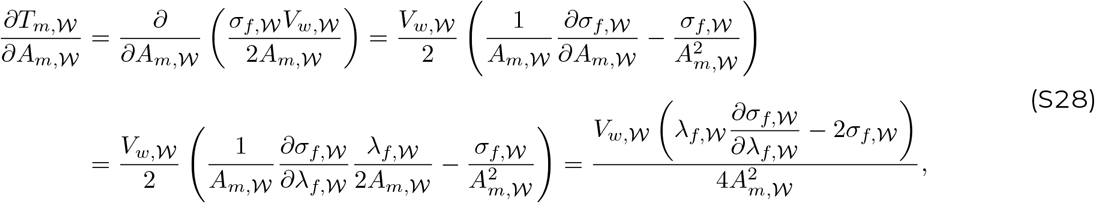

with the tangent modulus of the myofiber stress,

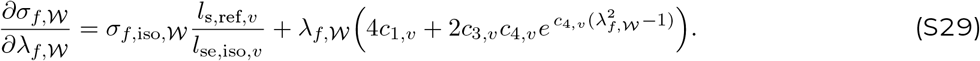

Equations (S28) and (S29) were evaluated at the predicted state, so that the slope in Eq. (S27) is constant throughout the corrector step.

##### Ventricular geometry

In the TriSeg formulation^10^, the ventricles are described as three spherical wall segments meeting at a common junction circle. The geometry is rotationally symmetric about the *x*-axis, which is normal to the plane of the junction circle. The *y*-axis lies in that plane. By convention, the *x*-axis is positive towards the right ventricle. Each wall segment W is a spherical cap. It is described by its height *x*_*m,W*_, measured from the junction plane to the apex, and by the radius *y*_*m*_ of its base circle, which is the junction circle shared by all three segments. The radius of curvature *r*_*m,W*_ and the enclosed volume *V*_*m,W*_ follow from *x*_*m,W*_ and *y*_*m*_,

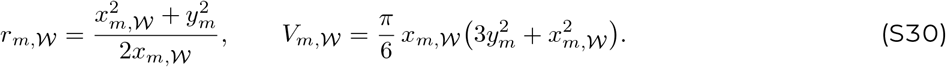

The ventricular geometry is therefore described by four parameters: the three cap heights *x*_*m,W*_ and the junction circle radius *y*_*m*_. The left free wall bulges away from the right ventricle and has *x*_*m*,LFW_ *<* 0. The septal and right free wall normally have *x*_*m, W*_ *>* 0. Four conditions determine these parameters: two imposed by the ventricular cavity volumes, and two by the equilibrium of the junction circle. The midwall of each ventricle encloses its cavity together with half of the volume of the two wall segments bounding it. The cavity volumes *V*_lv_ and *V*_rv_ follow from Eq. (S5). The wall volumes were obtained from the thin-wall approximation *V*_*w, W*_ = *A*_*m*,ref, *W*_ *W*_th, *W*_, with the midwall reference area and the wall thickness given in Table 2. The midwall volumes of the left and right ventricles are then

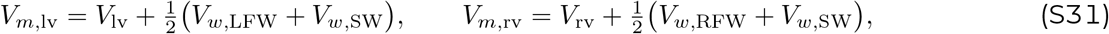

and geometrically the same volumes are the regions enclosed between neighbouring caps, which gives the first two conditions

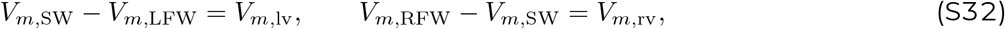

with each cap volume given by Eq. (S30) as a function of *x*_*m,W*_ and *y*_*m*_. The junction circle was assumed to carry no load of its own, so that the three midwall tensions acting on it must be in equilibrium. Each wall segment pulls on the rim in the meridional direction of its own surface, with a force per unit length equal to its midwall tension *T*_*m,W*_ . This pull is tangent to the cap and therefore perpendicular to the radius from the centre of curvature to the rim, which has the components *r*_*m,W*_ − *x*_*m,W*_ along the *x*-axis and *y*_*m*_ along the *y*-axis. Resolving the pull into a component *T*_*x,W*_ normal to the junction plane and a component *T*_*y,W*_ lying within it gives

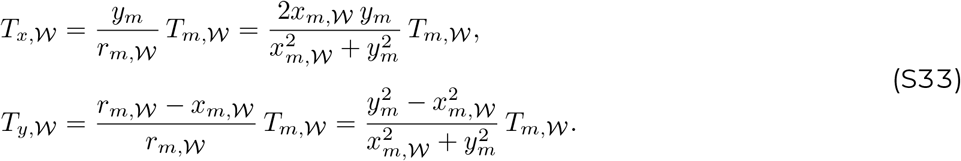

The remaining two conditions are that each component vanishes when summed over the three wall segments,

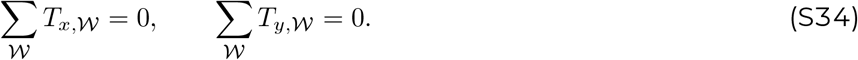

##### Numerical solution

The two volume conditions of Eq. (S32) were enforced exactly. At each time step, a Newton–Raphson scheme adjusted (*x*_*m*,SW_, *y*_*m*_) until ∥(*R*_*x*_, *R*_*y*_)∥_2_ *<* 10^*−*3^, with the residuals

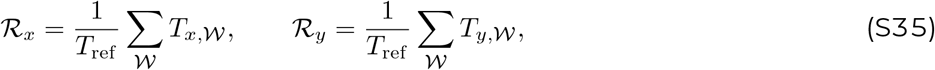

and 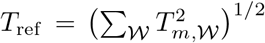. The scheme was initialised at the converged solution of the previous time step.

##### Recovered myofiber mechanics

At the converged geometry, the midwall area of each wall segment *A*_*m,W*_ follows from Eq. (S27). The corrected myofiber stretch is then 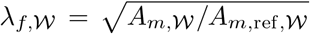, and the myofiber stress is re-evaluated from Eq. (S24) with *λ*_*f,W*_ in place of 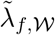 .

